# Structural and mechanistic insights into *Plasmodium* eIF2alpha dephosphorylation by UIS2 during the erythrocytic stage

**DOI:** 10.1101/2024.06.25.600622

**Authors:** Su Wu, Gerhard Wagner

## Abstract

UIS2 is a promising therapeutic target for malaria due to its essential role in the transmission and infectivity of *Plasmodium* to the host hepatocytes. In *Plasmodium*, UIS2 acts as a phosphatase toward phosphorylated eIF2alpha at Ser59, regulating translation initiation. However, its role during the erythrocytic stage, which causes clinical symptoms, remains unclear, and its protein structure for elucidating the dephosphorylation mechanism is unavailable. In this study, we analyzed the *Plasmodium* phenotype screening database and found that *UIS2*-deficient *Plasmodium* fails to proliferate during the erythrocytic stages. Single-cell transcriptomic data from the Malaria Cell Atlas revealed *UIS2* expression during both hepatic and erythrocytic stages, with significant upregulation in trophozoites and late schizonts, suggesting UIS2’s essential role in erythrocytic stage development. Structural analyses using AlphaFold modeling demonstrated that the UIS2 phosphatase domain (UIS2-PD) shares homology with human purple acid phosphatase, featuring conserved catalytic site residues. We observed that the N-terminal domain of UIS2 interacts with the S1 domain loop of a phospho-mimic mutant (Ser59Asp) eIF2alpha via electrostatic interactions. This protein interaction is distal to the phosphatase domain, indicating a unique substrate recruitment mechanism distinct from mammalian serine/threonine phosphatase PP1. Additionally, we identified a negatively charged cavity in UIS2-PD capable of binding the Ser59-containing loop in eIF2alpha for substrate recognition. Molecular docking studies showed that the phosphatase inhibitor salubrinal binds to this negatively charged cavity, potentially obstructing enzyme-substrate interactions. These structural insights into UIS2 and its interaction with eIF2alpha provide promising avenues for developing novel antimalarial drugs.

**Significance Statement:** Malaria poses a significant global health challenge, particularly with rising resistance to artemisinin therapies. This study reveals the critical role of UIS2 in *Plasmodium* proliferation during the symptomatic blood stage through phenotype screening and single-cell transcriptomics. We characterized the catalytic site within UIS2-PD and discovered a surface cavity for recognizing phosphorylated eIF2alpha by UIS2-PD, which can be inhibited by small molecules. Our findings identify UIS2 as a promising target for new antimalarial drugs, addressing both blood-stage parasites to alleviate clinical symptoms and liver-stage parasites for prophylactic treatment. This dual targeting strategy could prevent malaria onset post-exposure and reduce the overall parasite burden. The protein interaction between UIS2 and eIF2alpha is chemically targetable, facilitating the development of effective UIS2 inhibitors.

## Introduction

Malaria, caused by *Plasmodium* parasites, is a significant global health challenge with 249 million cases and 608,000 deaths reported in 2022 (1). *Plasmodium* parasites are transmitted through bites from infected *Anopheles* mosquitoes, with sporozoites migrating from mosquito salivary glands to the human body (2). After entry, sporozoites first invade liver cells, replicate, and release a small number of merozoites into the bloodstream (Fig. 1*A*). Merozoites then infect red blood cells (erythrocytes), progressing through ring, trophozoite, and schizont stages, ultimately causing cell rupture and new infection cycles. Clinical symptoms of malaria primarily arise from erythrocytic-stage parasites. Despite artemisinin-based combination therapies (ACTs) being the frontline treatment, the emergence of resistance poses a significant challenge to their efficacy (3). Thus, there is an urgent need for novel antimalarial targets and therapeutics.

**Figure 1.**
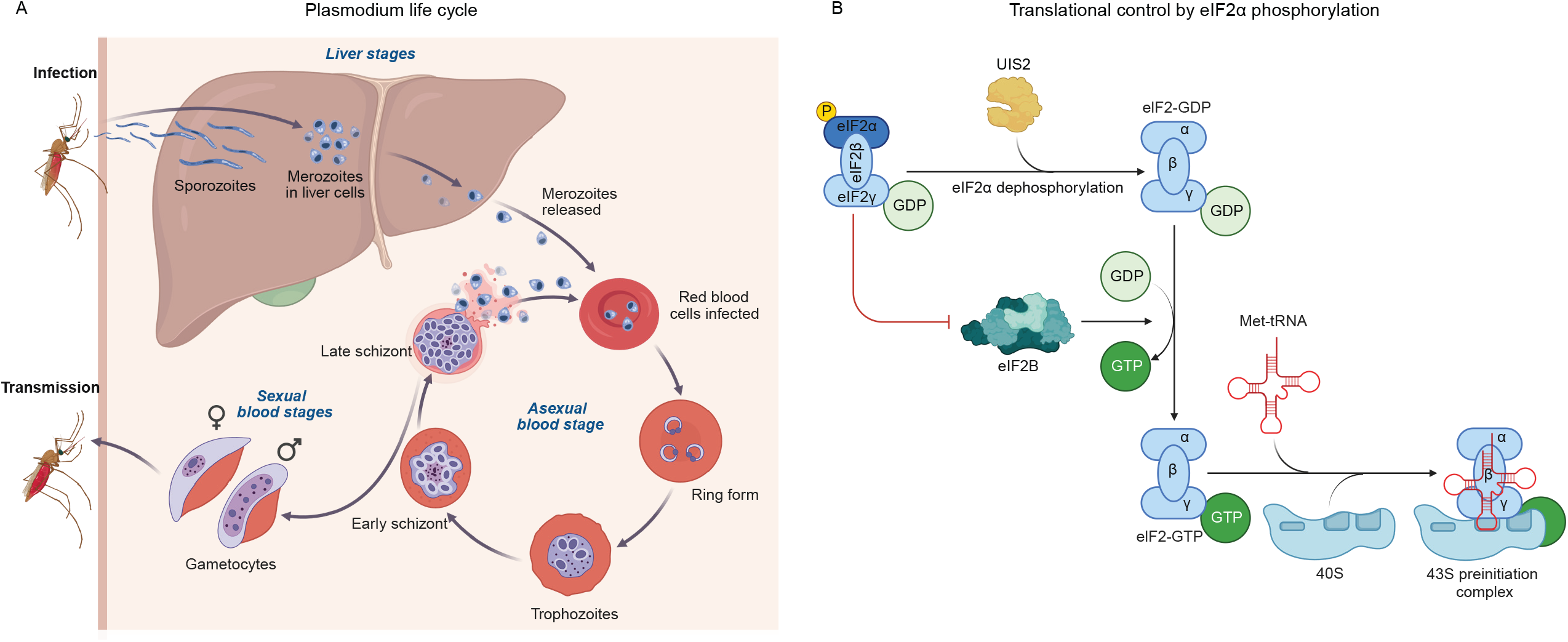
(*A*) The *Plasmodium* parasite life cycle in the human host involves hepatic and erythrocytic stages. During a blood meal, an infected *Anopheles* mosquito inoculates sporozoites into the human host. These sporozoites infect liver cells, mature, and then release merozoites into the bloodstream. Following this initial liver stage, the parasites undergo asexual multiplication stage in erythrocytes. Merozoites infect red blood cells, where they mature from the ring stage to trophozoites and then to schizonts, which rupture to release more merozoites. Some parasites differentiate into sexual erythrocytic stages, known as gametocytes, which can be transmitted back to the mosquito host. (*B*) UIS2 dephosphorylates eIF2alpha, promoting translation initiation. The GTP exchange factor eIF2B converts inactive eIF2-GDP to active eIF2-GTP. eIF2-GTP, along with Met-tRNAi, assembles with the 40S ribosome to form the 43S pre-initiation complex for translation initiation. When eIF2alpha is phosphorylated, eIF2B’s function to convert eIF2-GDP to eIF2-GTP is inhibited, leading to the inhibition of translation initiation.

The *Plasmodium* UIS2 (Up-regulated in Infective Sporozoites 2) protein is a promising therapeutic target for malaria intervention. Initially identified as a highly upregulated gene in sporozoites within mosquito salivary glands, *UIS2* is essential for parasite development (4). Knockout studies indicate that *UIS2*-deficient sporozoites can invade liver cells but fail to progress to hepatic-stage parasites (5). In sporozoites, *UIS2* is transcriptionally upregulated but translationally repressed by the RNA-binding protein Puf2. This mechanism allows *UIS2* mRNA to accumulate while limiting protein synthesis in quiescent sporozoites. Upon transmission to human hosts, the repression is lifted, enabling UIS2 protein production and facilitating the transition of sporozoites into hepatic-stage parasites (6). Despite UIS2’s critical role in parasite development at the hepatic stage, its role during the erythrocytic stages that cause clinical symptoms remains unknown. Understanding the functions of UIS2 during the blood stages of *Plasmodium* development is crucial for evaluating whether targeting UIS2 can enhance the effectiveness of treatments across multiple stages of the parasite’s life cycle.

UIS2 exhibits phosphatase activity by modulating the phosphorylation status of *Plasmodium* eIF2alpha, thereby facilitating translation initiation needed for protein synthesis during parasite development (5). eIF2alpha is a subunit of the eIF2 trimer complex. When eIF2alpha is bound to GTP, the eIF2 complex associates with initiator methionyl-tRNA (Met-tRNAi), the ribosomal 40S subunit, and other components to form the 43S pre-initiation complex, which scans mRNA for the initiation codon (Fig. 1*B*). In *Plasmodium*, eIF2alpha can be phosphorylated at serine 59 (P-eIF2alpha) by three kinases under different conditions: eIK1, eIK2, and PK4 (7). eIK1 phosphorylates eIF2alpha in response to amino acid deprivation (8), PK4 targets eIF2alpha in erythrocytic schizonts and gametocytes (9), and eIK2 phosphorylates eIF2alpha within sporozoites inside mosquito salivary glands (10). P-eIF2alpha inhibits eIF2B’s ability to convert eIF2 to its active GTP-bound state and thus suppresses translation initiation (11). After transmission to the host, UIS2 dephosphorylates P-eIF2alpha, enabling the protein synthesis required for sporozoites to exit latency and transition to the liver stage (5). However, the elucidation of UIS2’s structure and catalytic mechanism remains incomplete, hindering the development of effective chemical inhibitors.

In this study, we elucidated the critical role of UIS2 in *Plasmodium* growth during the asexual blood stage through analysis of the *Plasmodium* phenotype screening database from PlasmoGEM. By examining single-cell transcriptomic profile data from the Malaria Cell Atlas, we found *UIS2* overexpression in both liver and blood stages. Using Alphafold2.3 for molecular modeling, we identified significant structural homology between the phosphatase domain (PD) of UIS2 and human purple acid phosphatase, with conserved amino acids at the active site. Our molecular modeling revealed the protein interaction interface between UIS2 and a phospho-mimic mutant of eIF2alpha, showing that UIS2-NTD facilitates recruitment of the Ser59-containing loop (S59-loop) in eIF2alpha. Additionally, we demonstrated that UIS2-PD has a negatively charged cavity capable of binding the S59-loop in eIF2alpha, a region susceptible to targeting by salubrinal.

## Results

### *UIS2* is essential for *Plasmodium* growth during the erythrocytic stage, showing overexpression in both hepatic and erythrocytic stages

We first assessed the importance of translation initiation genes in *Plasmodium berghei* (*P. berghei*), a rodent malaria parasite, that shares similar symptoms with the predominant human malaria species, *Plasmodium falciparum* (*P. falciparum*). Using an *in vivo* phenotype screening database from PlasmoGEM, we analyzed the growth rates of *P. berghei* mutants during asexual blood stages in mice, each mutant lacking one of 2,578 *Plasmodium* genes (12). We found *UIS2* and several translation initiation genes, including *eIF4G, eIF4A3, eIF3D*, and *eIF3I*, are essential for *P. berghei* growth (Fig. 2*A*). Consistently, a previous study has shown that targeted *UIS2*-disruption in *P. falciparum* led to parasite death in cultured human erythrocytes (13). We also found that knockout of eIF2alpha kinases *eIK2* and *PK4* resulted in slow growth, indicating their non-essential role during the erythrocytic stage. These results suggest *UIS2* is vital for parasite multiplication during the erythrocytic stage, beyond its role in transitioning sporozoites to hepatic stages.

**Figure 2.**
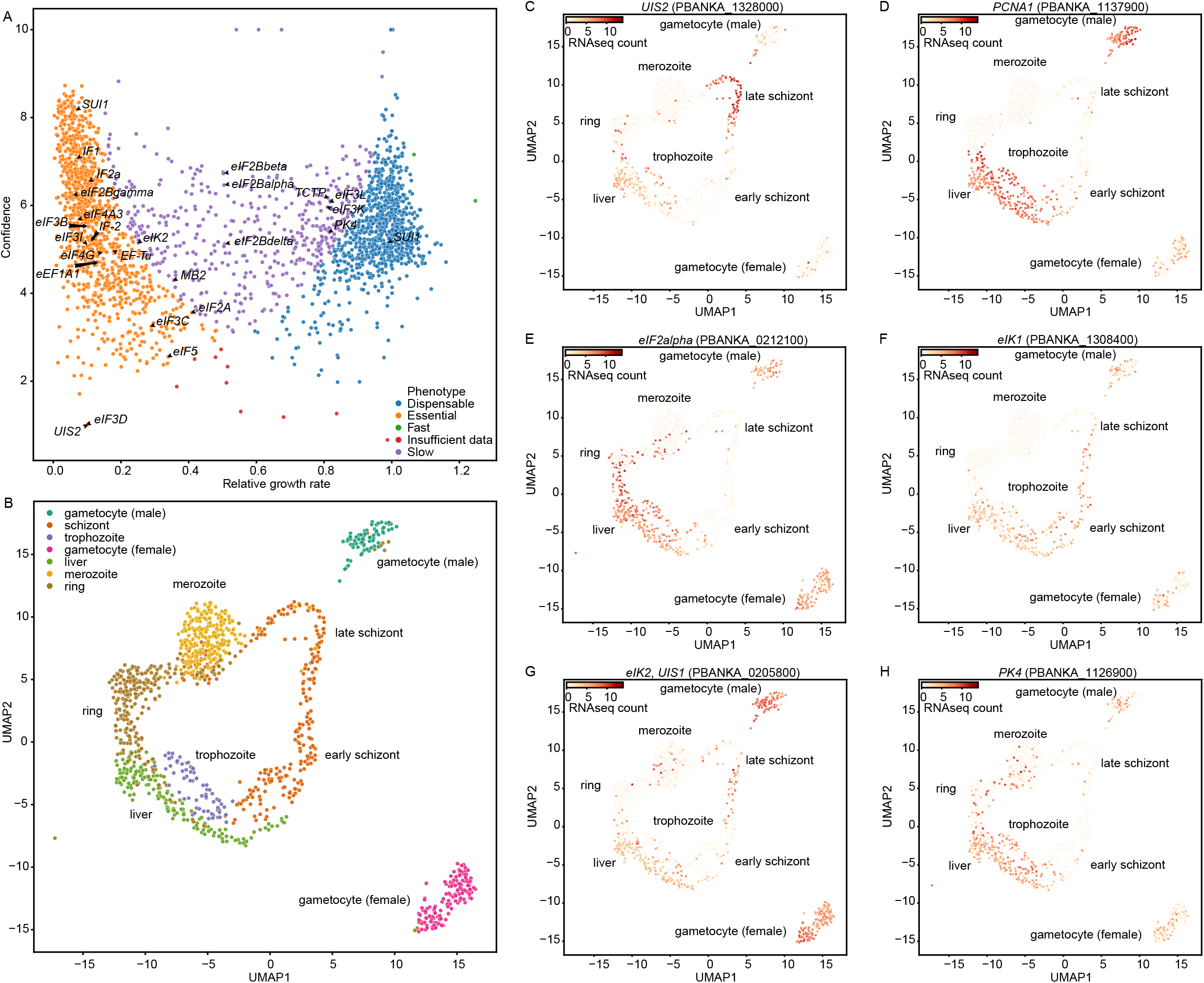
*UIS2* is essential for *Plasmodium* parasite growth in the erythrocytic stage, and overexpresses during the erythrocytic and hepatic stages. (*A*) The dot plot illustrates the relative growth rates of 2,578 mutants of *Plasmodium berghei*, each knocked out for individual parasite genes, following their pooled infection in mice as schizonts. Relative growth rates on the x-axis were determined by changes in the abundance of each barcoded knockout gene, normalized against the control post-infection, using next-generation sequencing. The variance in growth rate was calculated from the counts of the barcoded knockout gene across three replicate infections for each mutant. Confidence on the y-axis represents the negative logarithm of variance. The knockout genes were categorized and color-coded into four distinct growth phenotypes: essential, slow-growing, dispensable, and fast. Genes involved in translation initiation are specifically labeled. (*B*) The UMAP plot presents 1,137 individual *P. berghei* parasites across all life cycle stages within the mouse host, with parasites clustered based on transcriptome similarities. Single-cell RNAseq analysis was performed on parasites isolated from each life stage. Each parasite is color-coded according to its life stage. (*C*-*H*) The UMAP plots display the expression profiles of *UIS2* and its associated genes in the predominant life stage of parasites. Each life stage is labeled according to the clusters in (*B*). Parasites expressing the specified gene are colored red, with the intensity reflecting the relative expression levels, as determined by the RNA counts in the single-cell RNAseq dataset.

We analyzed *UIS2* expression throughout the parasite life cycle using the data from the Malaria Cell Atlas (14), which contains single-cell transcriptomic data for *P. berghei* parasites across various morphological stages within hosts (Fig. 2*B*). Elevated *UIS2* expression was observed during the hepatic stage and erythrocytic trophozoite stage in mouse host (Fig. 2*C*). These stages are critical for parasite proliferation, confirmed by co-expression of DNA replication markers *PCNA1* and *ORC1* (Fig. 2*D* and S1*A*). High *UIS2* expression was also observed in late schizonts, co-expressing with late schizont markers *CDPK5* and *HSP101* (15, 16) (Fig. S1 *B* and *C*). The late schizont stage involves cytokinesis of multinucleated schizonts, releasing merozoites and causing erythrocyte destruction, which leads to malaria symptoms such as fever and anemia. Since UIS2 protein translation is known to be repressed by Puf2 in sporozoites (5), we examined *Puf2* expression to understand its regulation in other *Plasmodium* stages. We found that *Puf2* strongly co-expressed with *UIS2* in mosquito sporozoites but minimally expressed in most parasite stages within the mouse host (Fig. S2 *A*-*D*). This suggests that UIS2 functions in both hepatic and erythrocytic stages due to transcriptional upregulation and the absence of Puf2 repression.

We also investigated mRNA expression of *eIF2alpha* and its kinases *eIK1, eIK2*, and *PK1*. Similar to *UIS2*, high expression levels were observed during the hepatic stage and in erythrocytic trophozoites (Fig. 2 *E*-*H*). Other translation initiation factors such as *eIF4E, eIF4G*, and *eIF3D* also overexpressed during these stages (Fig. S1 *D*-*F*), suggesting a high demand for translation initiation during parasite proliferation. Unlike the sporozoite stage, transcription and translation regulation for protein expression are tightly coupled in the erythrocytic stage, meaning that gene expression levels likely correlate with protein expression levels (17). Therefore, the elevated mRNA expression of *eIF2alpha* and its kinases in the erythrocytic stage, likely correspond to increased protein expression. These expression patterns suggest that UIS2 modulates eIF2alpha phosphorylation during proliferative stages, such as the hepatic stage and erythrocytic trophozoites, and contributes to the late schizont stage associated with malaria symptoms.

### The phosphatase domain of UIS2 shows structural homology with purple acid phosphatase

To understand the molecular mechanism of UIS2-mediated dephosphorylation on eIF2alpha in *Plasmodium*, we conducted structural analyses of the full-length UIS2 protein from *P. berghei* (*Pb*UIS2) using Alphafold2.3. *Pb*UIS2 comprises four domains: a short signal peptide (SP), an N-terminal helical domain (NTD), a phosphatase domain (PD) with an alpha-beta structure, and a C-terminal helical domain (CTD) (Fig. 3 *A* and *B*). Alphafold2.3 provides the predicted local distance difference test (pLDDT) to measure the local confidence of each residue in the structure. Most residues in UIS2-PD have pLDDT scores exceeding 90, indicating highly accurate predictions of UIS2-PD structure (Fig. S3 *A*-*C*). Furthermore, the predicted full-length UIS2 protein from *P. falciparum* (*Pf*UIS2) shares a highly similar overall structure with *Pb*UIS2 and exhibits high pLDDT scores in its phosphatase domain (Fig. S3 *D*-*F*).

**Figure 3.**
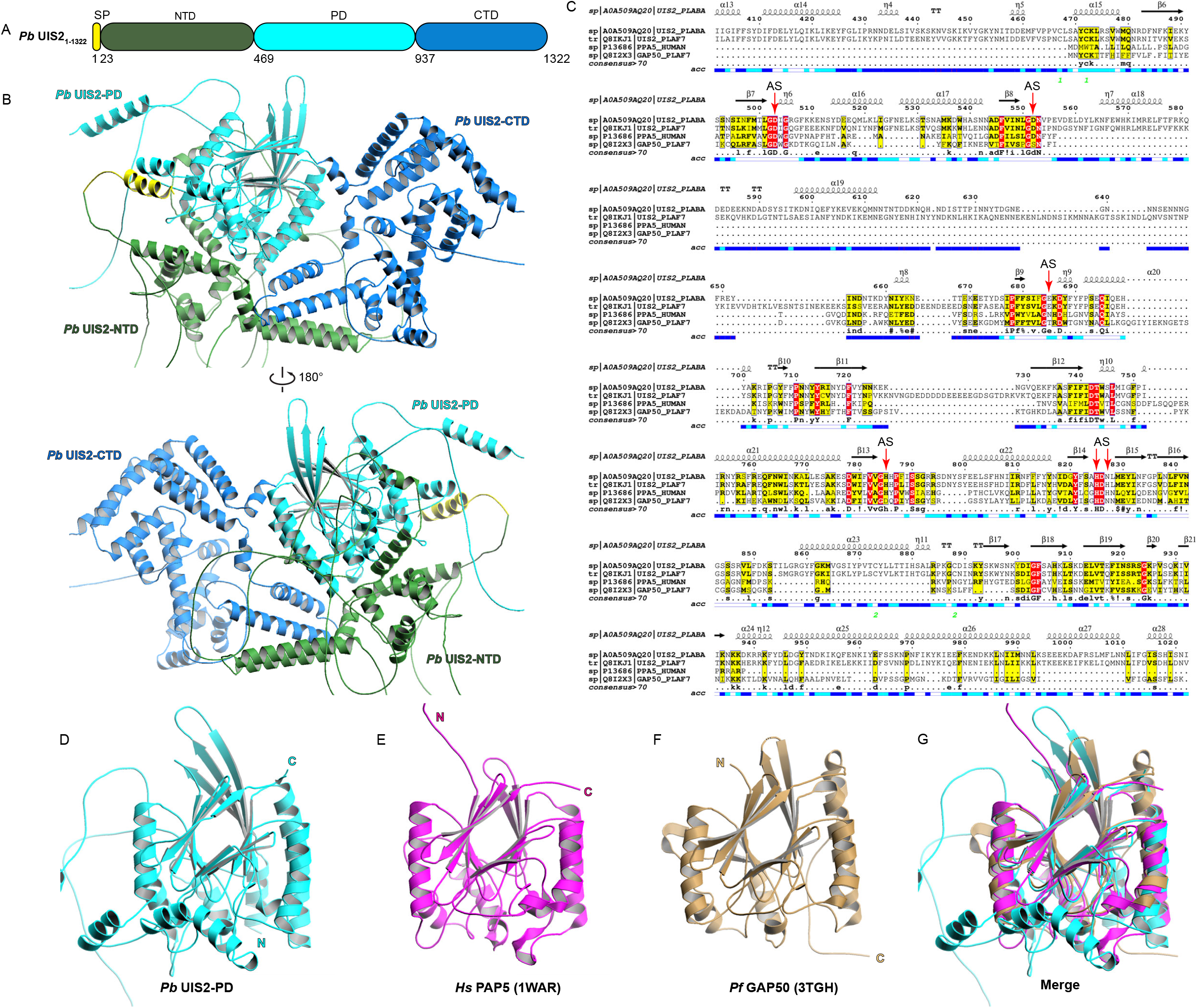
The phosphatase domain of UIS2 shows both sequence and structural homology with purple acid phosphatase. (*A*) The schematic diagram illustrates the domains within *Pb*UIS2: signal peptide (SP), N-terminal domain (NTD), phosphatase domain (PD), and C-terminal domain (CTD). (*B*) Predicted protein structure of full-length *Pb*UIS2 generated by Alphafold2. (*C*) Multiple sequence alignment of the UIS2 phosphatase domain (UIS2-PD) from *P. berghei* and *P. falciparum, Homo sapiens* PAP5, and *P. falciparum* GAP50. The alignment was performed using Clustal Omega and Escript 3.0. Secondary structure elements are indicated: α-helices (squiggles), 3_10_-helices (small squiggles), β-strands (arrows), β-turns (TT letters). Identical residues are highlighted in red boxes, and similar but significantly different residues are highlighted in yellow boxes. Accessibility is denoted by a bar below the alignment: blue for accessible, cyan for intermediate, and white for buried residues. Green digits indicate disulfide bridges at the bottom of sequence blocks. Active site residues are labeled “AS” and indicated with red arrows. (*D*) Visualization of the protein structure of the phosphatase domain in *Pb*UIS2 from (*B*). (*E*-*F*) Protein structures of *Hs*PAP5 and *Pf*GAP50, obtained from the Protein Data Bank. (*G*) Superposition of the phosphatase domain of *Pb*UIS2 with *Hs*PAP5 and *Pf*GAP50 structures.

Alphafold2.3 employed the HHsearch algorithm to identify structural homologs as templates for modeling UIS2. The top two structural homologs identified are *Homo sapiens* purple acid phosphatase (*Hs*PAP5, PDB ID: 1WAR) and the glideosome-associated protein GAP50 from *P. falciparum* (PDB ID: 3TGH). Multiple sequence alignment using Clustal Omega on *Pb*UIS2, *Pf*UIS2, *Hs*PAP5, and *Pf*GAP50 revealed many regions of sequence similarity within their phosphatase domains (Fig. 3*C*), supporting UIS2-PD, PAP5, and GAP50 as structural homologs. The predicted UIS2-PD structure, akin to the crystal structures of PAP5 and GAP50, features a central β-sandwich surrounded by α-helices on both sides (Fig. 3 *D*-*G*, and Fig. S4 *A*-*D*). *Pb*UIS2-PD contains an additional sequence stretch from residue 579 to 676 that lacks folded structure and appears as an extended loop. Both PAP5 and GAP50 belong to the superfamily of metallo-dependent phosphatases, which includes Ser/Thr protein phosphatases such as PP1. We thus compared the UIS2-PD structure with the human PP1/GADD35 structure (PDB ID: 4XPN) (Fig. S4 *E* and *F*). Although the catalytic domain of PP1 also comprises a central β-sandwich surrounded by α-helical domains (18, 19), the lengths of its β-sheets and α-helices are much shorter than those in UIS2-PD. In summary, these findings suggest a closer structural homology between UIS2-PD and PAP5 compared to its mammalian counterpart PP1.

### UIS2-PD contains highly conserved residues within its catalytic site

We conducted a comparative analysis of UIS2-PD with PAP5 and GAP50 to elucidate its structural features. UIS2-PD has a central β-sandwich consisting of two β-sheets with fourteen β-strands (β6 – β20), flanked by two α-helices (α17 and α21) and several smaller helical segments (Fig. 4*A*). The catalytic site is nestled within the β-sandwich, while an adjacent surface cavity likely serves as its entrance (Fig. 4*B*). PAP5 catalyzes the hydrolysis of phosphomonoesters and anhydrides under acidic conditions (20). Its catalytic site resides within the channel formed by the two β-sheets of its β-sandwich structure and contains a buried dinuclear metal center with Fe^2+^ and Fe^3+^, bridged by a phosphate and a water molecule (Fig. 4 *C* and *D*) (20, 21). The surrounding α-helices of UIS2-PD and PAP5 exhibit slightly different orientations for the β-sandwich. In contrast, GAP50 serves as a scaffold protein without enzymatic activity, functioning as an anchor for the glideosomal complex on the inner membrane of *Plasmodium* (22). Although the overall folds of GAP50 and PAP5 are similar, GAP50 lacks a dinuclear metal site (Fig. 4 *E* and *F*) (22). One Co^2+^ ion is located near the β-sandwich and exposed to the surface, while a second Co^2+^ ion is at the α-helix, away from the β-sandwich.

**Figure 4.**
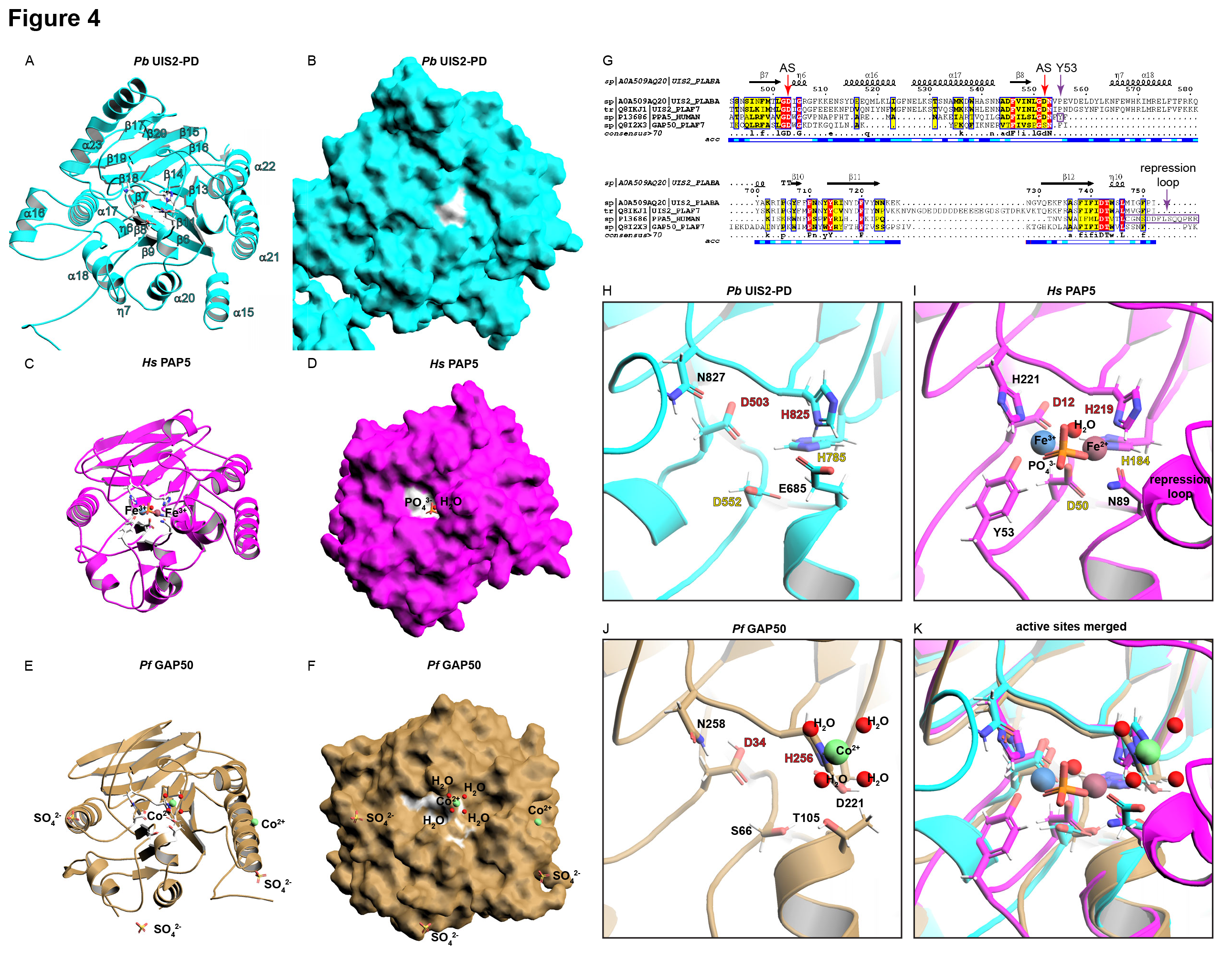
Structural comparison of the active sites of *Pb*UIS2-PD, *Hs*PAP5, and *Pf*GAP50. (*A*-*B*) The active site in the predicted phosphatase domain (PD) of *Pb*UIS2 is depicted, with predicted residues that form the catalytic pocket shown as sliver sticks. (*B*) Surface representation of (*A*). (*C*-*D*) The active site in *Hs*PAP5. (*D*) Surface representation of (*C*). (*E*-*F*) The active site in *Hs*GAP50. (*F*) Surface representation of (*E*). (*G*) Multiple sequence alignment of protein sequences reveals a unique Tyr53 and repression loop sequence in *Hs*PAP5, labeled in purple. Active site residues are labeled “AS” and indicated with red arrows. (*H*-*J*) Close-up views of the catalytic pockets in *Pb*UIS2-PD, *Hs*PAP5, and *Pf*GAP50, with the active site residues labeled. Identical active site residues shared among *Pb*UIS2-PD, *Hs*PAP5, and *Pf*GAP50 are highlighted in red. Identical active site residues shared specifically between *Pb*UIS2-PD and *Hs*PAP5 are highlighted in yellow. The phosphate ion is depicted in orange sticks. The ferric ion is represented as a blue sphere, the ferrous ion as a dark red sphere, and the cobalt ion as a green sphere. Water molecules are illustrated as red spheres. (*K*) Superposition of key residues at the active site from *Pb*UIS2-PD (cyan) with those of *Hs*PAP5 (magenta) and *Pf*GAP50 (brown).

We conducted a comparative analysis of the active sites of UIS2-PD and PAP5 to elucidate their catalytic mechanisms. In the catalytic center of PAP5, two iron atoms, Fe^2+^ and Fe^3+^, coordinate with a phosphate, a water molecule, three histidines (His184, His219, and His221), two aspartic acids (Asp12 and Asp50), and one asparagine (Asn89) (20) (Fig. 4*I*). In the catalytic center of UIS2-PD, we identified two conserved histidines (His825 and His785), and two conserved aspartic acids (Asp503 and Asp552) (Fig. 4*H*). Additionally, two less conserved residues from UIS2, Glu685 and Asn827, occupy positions analogous to residues Asn89 and His221 in PAP5’s catalytic center. In PAP5’s active site, the Tyr53 residue undergoes a charge-transfer transition with Fe^3+^, resulting in the enzyme’s purple color (23). However, Tyr53 is unique to PAP5, with no analogous residue found in UIS2-PD’s active site (Fig. 4*G*). Furthermore, there is a repression loop (Cys140 to Arg153) adjacent to the PAP5’s active site (Fig. 4*I*). This repression loop can be post-translationally cleaved to regulate PAP5’s enzymatic activity (24), a feature absent in UIS2-PD (Fig. 4*G*). In contrast, the supposed catalytic center of GAP50 differs from that of UIS2-PD and PAP5, with poorly conserved residues (Fig. 4 *J* and *K*). Only Asp34 and H256 are conserved in GAP50, with His256 coordinating a Co^2+^ ion along with four water molecules. This disparity explains the lack of enzymatic activity in GAP50. In summary, these results suggest that the catalytic centers of UIS2-PD and PAP5 are highly conserved, indicating potential conservation in their catalytic mechanisms. The primary differences between the active sites from UIS2-PD and PAP5 are the absence of a repression loop and the Tyr53 residue in UIS2-PD.

### Interaction between UIS2-NTD and the S59-loop in *Pb*eIF2alpha is distal to UIS2-PD

Previous biochemical mapping studies found that P-eIF2alpha, rather than its unphosphorylated counterpart, binds to *Pb*UIS2-NTD, indicating a regulatory role for UIS2-NTD in the dephosphorylation process (5). Initially, we used Alphafold2 to model *Pb*eIF2alpha and *Pf*eIF2alpha individually as monomers. *Pb*eIF2alpha comprises an N-terminal domain (NTD) containing the S1-type oligonucleotide/oligosaccharide binding-fold subdomain (S1 domain) and the α-helical subdomain, along with a C-terminal domain (Fig. 5 *A* and *B*). Both *Pb*eIF2alpha and *Pf*eIF2alpha share high sequence homology with human eIF2alpha, with most residues being identical throughout the sequence (Fig. S5*A*). Most residues in *Pb*eIF2alpha and *Pf*eIF2alpha have pLDDT scores above 70, and those in the S1 domain surpass 90, indicating highly accurate structure predictions (Fig. S5 *B* to *E*). The overall folds of *Pb*eIF2alpha and *Pf*eIF2alpha closely resemble the NMR and crystal structures of human eIF2alpha (Fig. S5 *F* and *G*) (25, 26).

**Figure 5.**
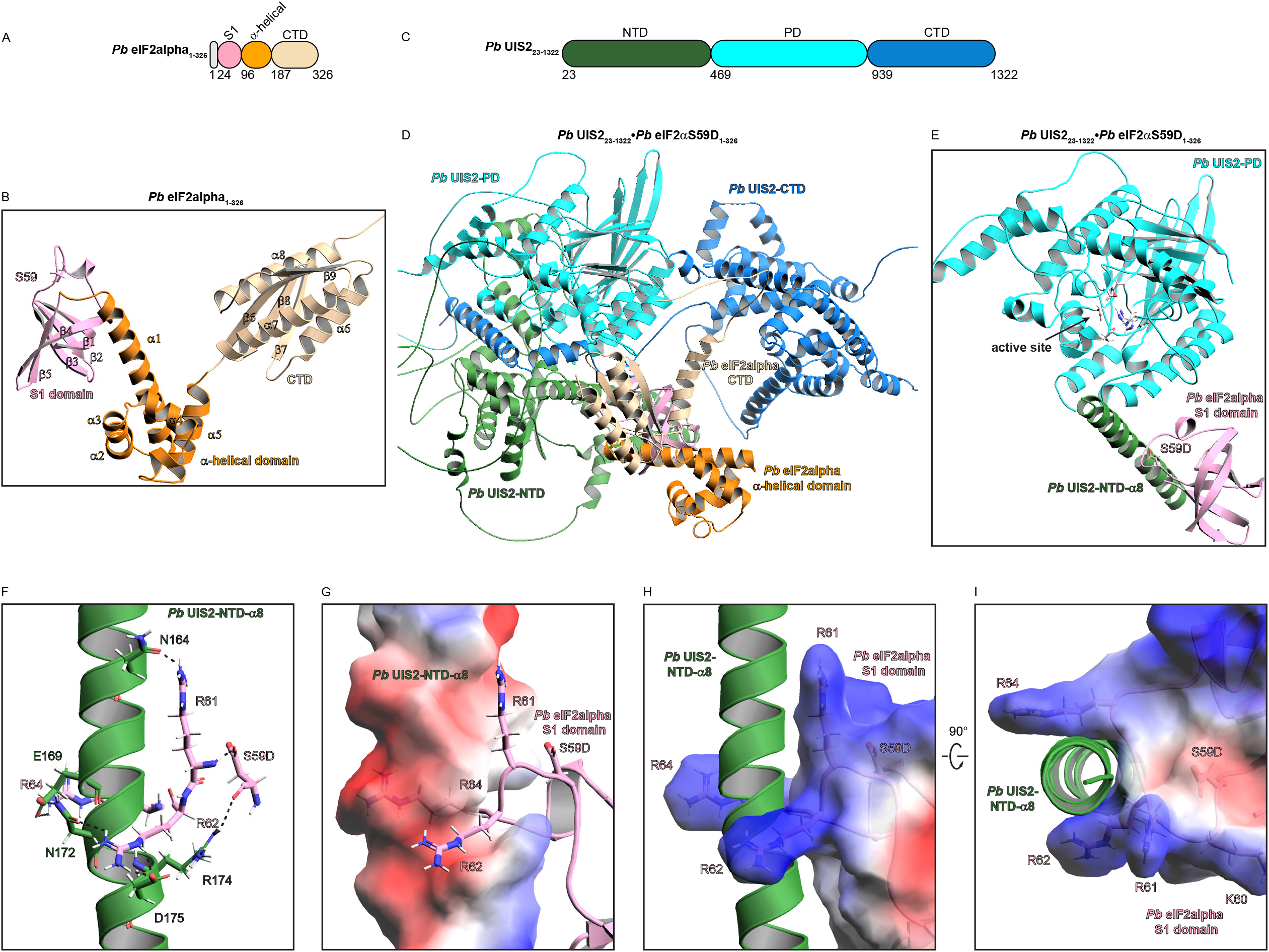
The N-terminal domain of *Pb*UIS2 recruits the S59-loop in *Pb*eIF2alpha. (*A*) The diagram illustrates the domains of eIF2alpha. (*B*) The cartoon depicts the structure of eIF2alpha from *P. berghei*, predicted by the AlphaFold2 monomer approach. (*C*) The diagram illustrates the *Pb*UIS2 construct used in the AlphaFold modeling. (*D*) The cartoon shows the protein complex structure formed by *Pb*UIS2 (residues 23-1322) and *Pb*eIF2alpha with Ser59 mutated into aspartic acid (S59D) to mimic phosphorylation, predicted by the AlphaFold2 multimer approach. (*E*) An isolated view of the *Pb*UIS2 (residues 23-1322) and *Pb*eIF2alphaS59D complex from (*D*), showing only the phosphatase domain (PD) and helix 8 (α8) from UIS2-NTD, along with the S1 domain of *Pb*eIF2alphaS59D. Key residues in the active site of *Pb*UIS2-PD are depicted as sliver sticks. (*F*) A close-up view of the interaction interface between *Pb*UIS2-NTD and the loop region from the S1 domain of *Pb*eIF2alphaS59D. Polar interactions are represented as black dashes. (*G*) The α8 helix from *Pb*UIS2-NTD is displayed as an electrostatic surface, with red indicating negative charges and blue indicating positive charges. The loop region from the *Pb*eIF2alphaS59D S1 domain is depicted as cartoons, with residues involved in protein interaction shown as sticks. (*H*-*I*) The structure of α8 helix from *Pb*UIS2-NTD is shown as a cartoon, along with the electrostatic surface of the loop region from the *Pb*eIF2alphaS59D S1 domain in two orientations.

We also modeled only the UIS2-NTD region using AlphaFold2 (Fig. S6*A*). However, Alphafold2 did not identify any homologous protein structure with HHsearch, and yielded generally low pLDDT scores (Fig. S6*B*), suggesting a lack of a defined structural motif in UIS2-NTD. However, two prominent α-helices, α8 and α9, are predicted to be present in the UIS2-NTD, with α8 featuring a negatively charged section (Fig. S6*C*).

To investigate the dephosphorylation mechanism of eIF2alpha by UIS2, we used Alphafold2 multimer approach to model the complex structure of *Pb*UIS2 bound to *Pb*eIF2alpha. Using a phospho-mimic mutant of *Pb*eIF2alpha, where the Ser59 residue was substituted by aspartic acid to emulate the phosphorylated state, alongside the UIS2_23-1322_ construct without the signal peptide (Fig. 5*C*), we generated the Alphafold-modeled protein complex. This modeled complex showed the interaction between the S1 domain of eIF2alpha and UIS2-NTD, consistent with experimental observations (Fig. 5 *D* and *E*) (5). Notably, the protein interaction interface was identified at α8 of the UIS2-NTD and the S59-loop of eIF2alpha, distal to the UIS2-PD (Fig. 4*E*). Due to the absence of a defined motif in UIS2-NTD and the S59-loop of eIF2alpha, the pLDDT scores for this interaction were low (Fig. S6 *D* and *E*). Nonetheless, the Alphafold model indicated that the eIFalpha2 Ser59Asp residue interacts with Arg174 from α8 of UIS2-NTD through electrostatic interaction (Fig. 5*F*). Specifically, three positively-charged arginines within the S59-loop in eIF2alpha formed a finger-like structure to anchor the loop onto the negatively-charged α8 of UIS2-NTD (Fig. 5 *G* to *I*).

Additionally, we employed Alphafold2 to model the complex structure of *Pf*UIS2 bound to *Pf*eIF2alphaSer59Asp. The resulting complex closely resembled the complex structure of *Pb*UIS2 and *Pb*eIF2alphaSer59Asp (Fig. S7 *A* and *B*) with a similar interaction predicted between the S59-loop and the α8 of *Pf*UIS2-NTD (Fig. S7*C*). Again, the interaction was facilitated by arginines from the S59-containing loop and α8 from *Pf*UIS2-NTD through electrostatic interaction (Fig. S7 *D* and *E*), suggesting a resemblance in UIS2 and eIF2alpha interaction mechanisms between *P. berghei* and *P. falciparum*. In summary, these AlphaFold2-modeled UIS2 and eIF2alpha protein complexes recapitulate the biochemical interactions observed in experimental results and suggest that UIS2-NTD likely recruits P-eIF2alpha through electrostatic interaction.

### The *Pb*UIS2-PD harbors a negatively charged surface cavity for substrate and salubrinal binding

The modeled complex of *Pb*UIS2 and *Pb*eIF2alphaSer59Asp reveals an interaction site between UIS2-NTD and eIF2alpha, situated distally from the UIS2-PD domain (Fig. 5*E*). This prompts the question of how UIS2-PD accesses its substrate, phosphorylated Ser59 from eIF2alpha, to catalyze the hydrolysis reaction. To elucidate the substrate recognition mechanism of UIS2-PD, we analyzed the surface map of UIS2-PD in the full-length UIS2 structure, modeled using the Alphafold2 monomer approach. We identified a surface cavity adjacent to the active site within the β-sandwich structure (Fig. 6*A*). Utilizing Consurf, we evaluated the evolutionary conservation of amino acid residues in UIS2, mapping the conservation scores onto the protein structure. Notably, this surface cavity in UIS2-PD appears to be a highly conserved region (Fig. 6*B*), characterized by negative surface charges (Fig. 6*C*) attributed to Asp503, Asp552, and Glu685 residues from the active sites (Fig. 4*H*). We hypothesize that following recruitment by UIS2-NTD, P-eIF2alpha may interact with this surface cavity of UIS2-PD as a substrate before reaching the catalytic center for hydrolysis.

**Figure 6.**
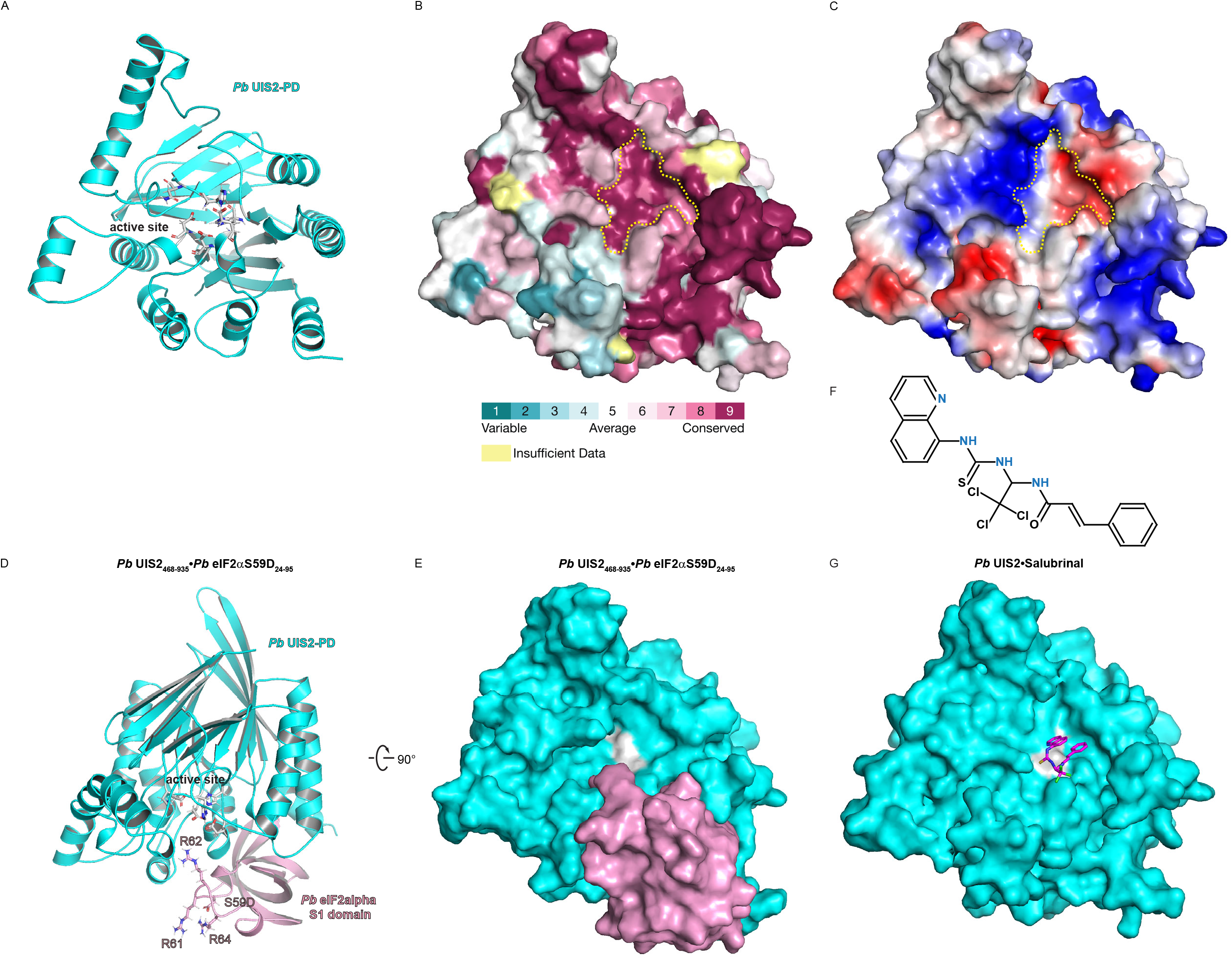
The active site of *Pb*UIS2-PD harbors a negatively charged cavity for substrate and salubrinal binding. (*A*) The protein structure of full-length *Pb*UIS2 was predicted by Alphafold2, with *Pb*UIS2-PD shown as a cartoon with the active site residues depicted as sliver sticks. (*B*) ConSurf analysis of *Pb*UIS2-PD, with the conserved catalytic cavity delineated by a yellow dashed line. (*C*) The electrostatic surface of *Pb*UIS2-PD. (*D*) The protein complex structure of *Pb*UIS2 (residues 468-935) and *Pb*eIF2alpha S59D (residues 24-95), as predicted by the AlphaFold2 multimer approach, is shown as cartoons. (*E*) Surface representation of the protein complex in (*D*). (*F*) The chemical structure of salubrinal is depicted, highlighting the nitrogen atom of the quinoline ring and the secondary amine groups in blue. (*G*) Docking of salubrinal into the catalytic cavity of *Pb*UIS2-PD using AutoDock Vina.

To investigate the substrate binding capability of UIS2-PD, we utilized the Alphafold2 multimer approach to predict the complex structure of UIS2-PD with the S1 domain of eIF2alpha, where Ser59 was mutated to aspartic acid. The resulting Alphafold-modeled structure reveals the positioning of the S59-containing loop at the surface cavity of UIS2-PD (Fig. 6 *D* and *E*). This result suggests that the interaction between the positively charged Arg62 residue and the negatively charged surface cavity may contribute to substrate binding or specificity. Furthermore, salubrinal (Fig. 6*F*) has been shown to inhibit the phosphatase activity of UIS2, leading to enhanced P-eIF2alpha in *Plasmodium* sporozoites (5, 27). We conducted molecular docking of salubrinal onto the Alphafold-predicted full-length UIS2 protein structure, using Autodock Vina. The top-scored docking positions of salubrinal were within UIS2-PD’s surface cavity (Fig. 6*G*). No chemical bond formation was evident between salubrinal and UIS2-PD in the docking model. This lack of bond formation might be attributed to salubrinal’s lesser potency as a small molecule inhibitor for UIS2 because salubrinal inhibits UIS2’s phosphatase activity only at very high concentrations (50 μM) (5). However, salubrinal comprises a quinoline and three secondary amine groups situated between its two ring structures (Fig. 6*F*), and these groups bear positive charges at physiological pH (28), which might explain salubrinal’s affinity for the negatively charged surface cavity within UIS2-PD. Furthermore, the positioning of salubrinal within UIS2-PD’s surface cavity implies its potential role in obstructing enzyme-substrate binding.

## Discussion

### The critical roles of UIS2 in the erythrocytic stage of *Plasmodium*

Our study highlights the essential role of UIS2 during the erythrocytic stages of *Plasmodium. UIS2*-knockout significantly impedes *Plasmodium* proliferation during the erythrocytic stage (Fig. 2*A*), suggesting its critical function beyond the hepatic stage. We observed robust *UIS2* expression in trophozoites, which are characterized by extensive DNA replication and metabolic activities, including multiple rounds of nuclear division (29, 30). Previous studies have shown the absence of P-eIF2alpha in *Plasmodium* trophozoites (27). Furthermore, the co-expression of *UIS2, PK4*, and *eIF2alpha* at the trophozoite stage (Fig. 2 *C, E* and *H*) suggests that UIS2’s role in dephosphorylating eIF2alpha is predominant over the kinase activity of PK4 in phosphorylating eIF2alpha. Additionally, *UIS2* expression was strong in late schizonts, the stage that undergoes cytokinesis, releasing daughter merozoites and causing erythrocyte rupture (31). During the late schizont stage, the mRNA levels of *eIF2alpha* and its kinases are low, and previous studies have observed P-eIF2alpha at the late schizont stage(27). This suggests that UIS2 may have other substrates besides eIF2alpha during the late schizont stage. Targeting UIS2 could thus offer dual advantages: mitigating the clinical symptoms caused by blood-stage parasites and providing prophylactic treatment against liver-stage parasites, thereby presenting a comprehensive therapeutic strategy.

### UIS2-NTD recruits the S59-loop in eIF2alpha

While prior biochemical studies have categorized UIS2 as a serine/threonine phosphatase with specificity towards P-eIF2alpha (5, 32), our research shows a structural similarity between UIS2-PD and human PAP5. Purple acid phosphatase and PP1 use different mechanisms for substrate recognition. While purple acid phosphatase uses residues around the active sites for substrate specificity (33), mammalian serine/threonine phosphatases often require an additional regulatory domain (34).

Previous biochemical studies suggested that UIS2-NTD functions as a regulatory domain for UIS2-PD (5). The Alphafold model recapitulated the interaction between UIS2-NTD and the S59-loop in eIF2alpha, but this interaction occurs distally to the UIS2-PD domain. This differs from PP1’s regulatory process, where the regulatory subunit like GADD34 binds directly to the PP1 catalytic domain to modify the catalytic cleft for substrate binding (18, 35). The S59-loop interacts with both UIS2-NTD and the surface cavity in UIS2-PD via the Arg residues, implying that the substrate is first recruited by UIS2-NTD before being recognized by UIS2-PD for catalysis.

This recruitment by UIS2-NTD may serve two purposes. First, UIS2 is a membrane protein located at the parasitophorous vacuole membrane (PVM), a barrier between the parasite and the host cell cytoplasm, during the blood stages of *P. falciparum* (13). As the membrane-targeting motif lies within UIS2-NTD (36), UIS2-NTD likely recruits P-eIF2alpha to the PVM for dephosphorylation. Second, P-eIF2alpha typically forms a nonproductive complex with eIF2B, where the phosphorylated Serine-containing loop binds to the cavity between eIF2Balpha and eIF2Bdelta, inhibiting the eIF2B complex’s ability to exchange GTP (37-39). UIS2-NTD may facilitate more efficient competition with eIF2B for P-eIF2alpha binding before initiating its phosphatase activity on UIS2-PD.

### The pocket in UIS2-PD recognizing the S59-loop in eIF2alpha is chemically targetable

Our results suggest that UIS2-PD may employ a substrate recognition mechanism similar to that of purple acid phosphatase. The positively charged S59-loop in eIF2alpha binds to a negatively charged surface cavity proximal to the catalytic site in UIS2-PD, indicating potential substrate recognition through electrostatic interaction. Although previous biochemical studies did not show UIS2-PD pulling down P-eIF2alpha *in vitro* (5), it is possible that eIF2alpha was rapidly dephosphorylated and released. Additionally, UIS2-PD lacks the repression loop found in PAP5’s catalytic site, which typically restricts substrate access to the active site (20, 40). This absence may confer UIS2-PD a lower Km and a faster substrate turnover rate compared to PAP5, although further experimental investigation is needed to verify this possibility.

Our findings also suggest that salubrinal competes with the S59-loop for binding to the surface cavity within UIS2-PD, thereby inhibiting protein-protein interaction. This aligns with a previous molecular docking study showing that salubrinal can fit into the catalytic pocket of GADD34/PP1 (41). Moreover, salubrinal inhibits P-eIF2alpha dephosphorylation in human cells by disrupting the formation of the GADD34/PP1 complex (42). We propose that UIS2-PD governs substrate specificity and is a viable target for chemical intervention.

In conclusion, our study reveals the crucial role of UIS2 in *Plasmodium* development, particularly during the erythrocytic stages. Targeting UIS2 offers the potential for developing treatments that address both hepatic and erythrocytic stages of malaria. Further research on UIS2 mechanisms and inhibitors could lead to effective antimalarial therapies.

## Materials and Methods

### Essential gene analysis and single-cell gene expression analysis

The phenotype screening data were sourced from PlasmoGEM (12). To create the phenotype plot, the scatterplot() function from the Python package “seaborn” was used, with translation initiation-related genes annotated from the phenotype dataset. Single-cell transcriptome profiles of 1,787 cells covering all life cycle stages were obtained from the Malaria Cell Atlas (dataset name: P. berghei SS2 set1). Stage annotation data were extracted from “pb-ss2-set1-ss2-data.csv,” and single-cell RNA-seq data from “pb-ss2-set1-ss2-exp.csv.” UMAP coordinates for each cell, provided in the original publication (14), were used as given in the “pb-ss2-set1-ss2-data.csv” file. Cells were colored based on gene expression values from the “pb-ss2-set1-ss2-exp.csv” file. Data selection and merging were performed using the Python package “pandas,” and plots were generated using the scatterplot() function from the “seaborn” package.

### Protein structure modeling

Protein structure predictions were performed using AlphaFold software version 2.3.1, an extension of the AlphaFold2 algorithm capable of predicting protein complexes as single entities (43). All tasks were executed on the high-performance computing (HPC) cluster at Harvard Medical School. Hardware varied based on availability, utilizing either NVIDIA A100 (80GB VRAM) or NVIDIA RTX8000 (48GB VRAM) GPUs, with an 80GB RAM reservation for each job. The “full_dbs” setting was used for the “db_preset.” Amino acid sequences were obtained from the NCBI database with the following accession numbers: *Pb*UIS2 (A0A509AQ20), *Pf*UIS2 (XP_001348788), *Pb*eIF2alpha (A0A509AJP9), and *Pf*eIF2alpha (Q8IBH7). The “monomer” model preset was used for single protein structure predictions, while the “multimer” model preset was employed for predicting protein complexes. The consistency of all prediction models generated was evaluated, and the top-ranked model for each task was selected for illustration using PyMOL. All non-Alphafold calculations were conducted on a System76 “Serval” mobile workstation with an Intel Core i7-8700k, running Pop!_OS 22.04 LTS (44).

### Multiple sequence alignment

Clustal Omega was used for new multiple sequence alignment with the protein sequences with the following accession numbers: *Pb*UIS2 (A0A509AQ20), *Pf*UIS2 (Q8IKJ1), *Hs*PAP5 (P13686), *Pf*GAP50 (Q8I2X3), *Pb*eIF2alpha (A0A509AJP9), *Pf*eIF2alpha (Q8IBH7) and *Hs*eIF2alpha (P05198). The aligned sequence results were visualized using the ESPript3 program.

### Molecular docking assay

Molecular docking studies were conducted using AutoDock Vina to predict the binding interactions of salubrinal with the phosphatase domain of *Pb*UIS2 (*Pb*UIS2-PD). The full-length UIS2 protein structure, predicted by AlphaFold2, was prepared using AutoDockTools by adding polar hydrogens, and merging non-polar hydrogens. Salubrinal structure was downloaded from the Zinc database and converted to PDBQT format with OpenBabel. The docking grid covers the entire *Pb*UIS2-PD. Docking simulations were performed with an exhaustiveness of 32. Docking results were evaluated based on binding affinity scores. The best-scoring pose was visualized using PyMOL to examine interactions between salubrinal and active site residues.

## Acknowledgments

This work was supported by the National Institute of Allergy and Infectious Diseases (5P01AI143565-03, to G.W.). We thank Drs. Philipp Aschauer and Christoph Gorgulla for their valuable scientific discussions.

