## Supplemental information for "Structural and mechanistic insights into *Plasmodium* eIF2alpha dephosphorylation by UIS2 during the erythrocytic stage"

**This PDF file includes:**

Figures S1 to S7

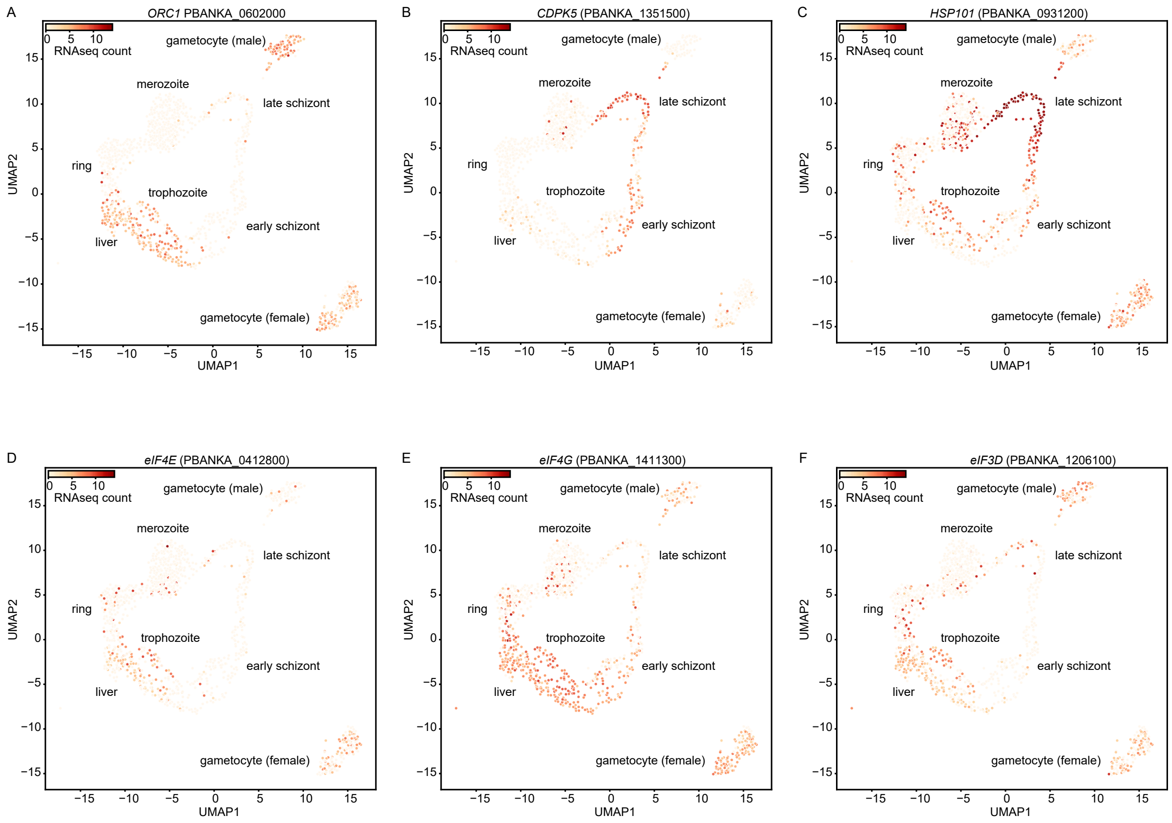


**Figure S1.** Expression of eIF4F complex components primarily occurs in the liver and trophozoite stages of parasites within the mouse host. (*A*-*C*) UMAP plots illustrate the expression profiles of marker genes specifically expressed in the liver, trophozoite, and late schizont stages of parasites. (*D*-*F*) UMAP plots depict the expression profiles of genes encoding the eIF4F complex components.

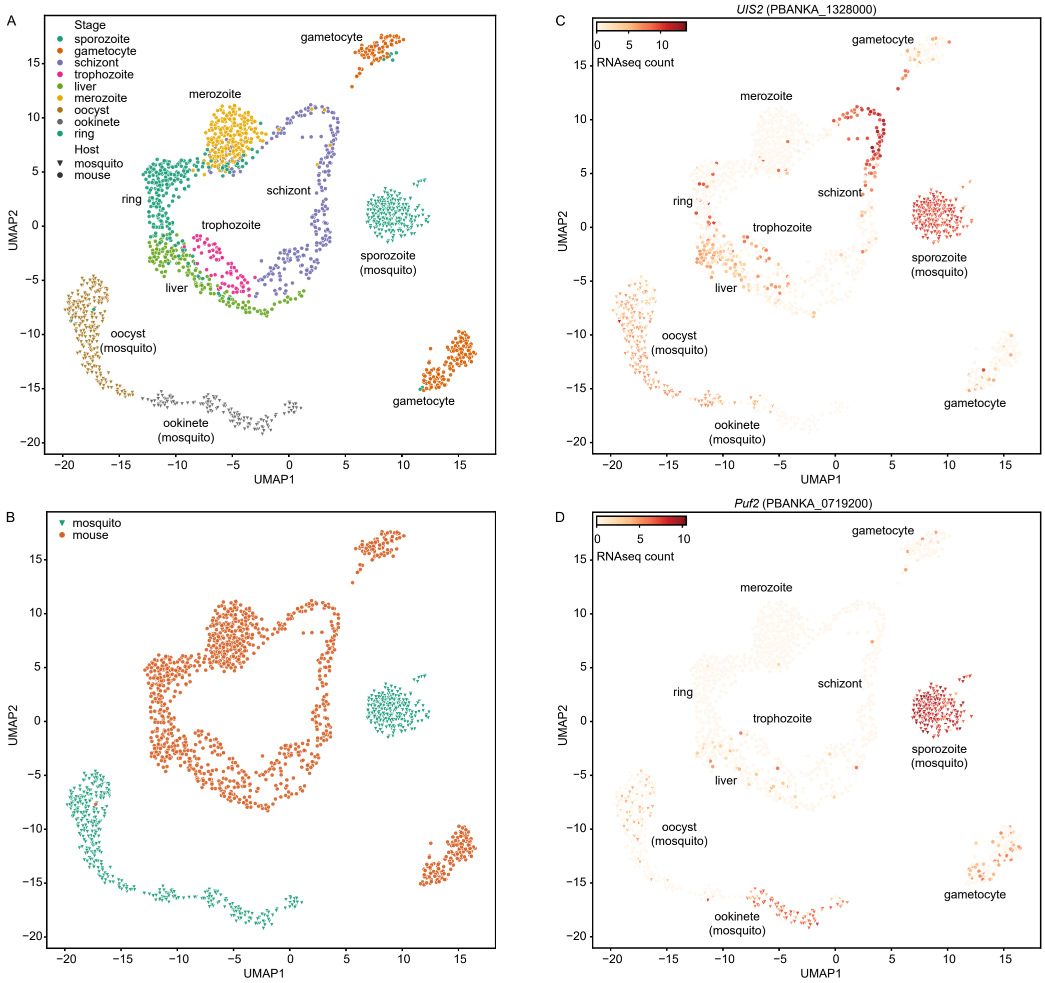


**Figure S2.** Translation repression of *UIS2*, primarily occurs in the parasite stages within the mosquito host. (*A*) UMAP plot displaying 1,787 individual *P. berghei* parasites across all life cycle stages, clustered based on transcriptome similarity. Parasites from each life stage were experimentally isolated and pooled for single-cell RNAseq analysis. Each parasite is colored according to its specific life stage. (*B*) UMAP plot as in (*A*) is colored based on the profile of the mouse or mosquito hosts from which individual *P. berghei* parasites were isolated. (*C-D*) UMAP plots depict the expression profiles of *UIS2* and *Puf2* in the life stages of parasites. Each life stage is labeled according to the clusters in (*A*) and (*B*). Cells expressing the indicated gene are shown in red, with the intensity of color reflecting the relative expression levels, determined by the RNA counts in the single-cell RNAseq dataset.

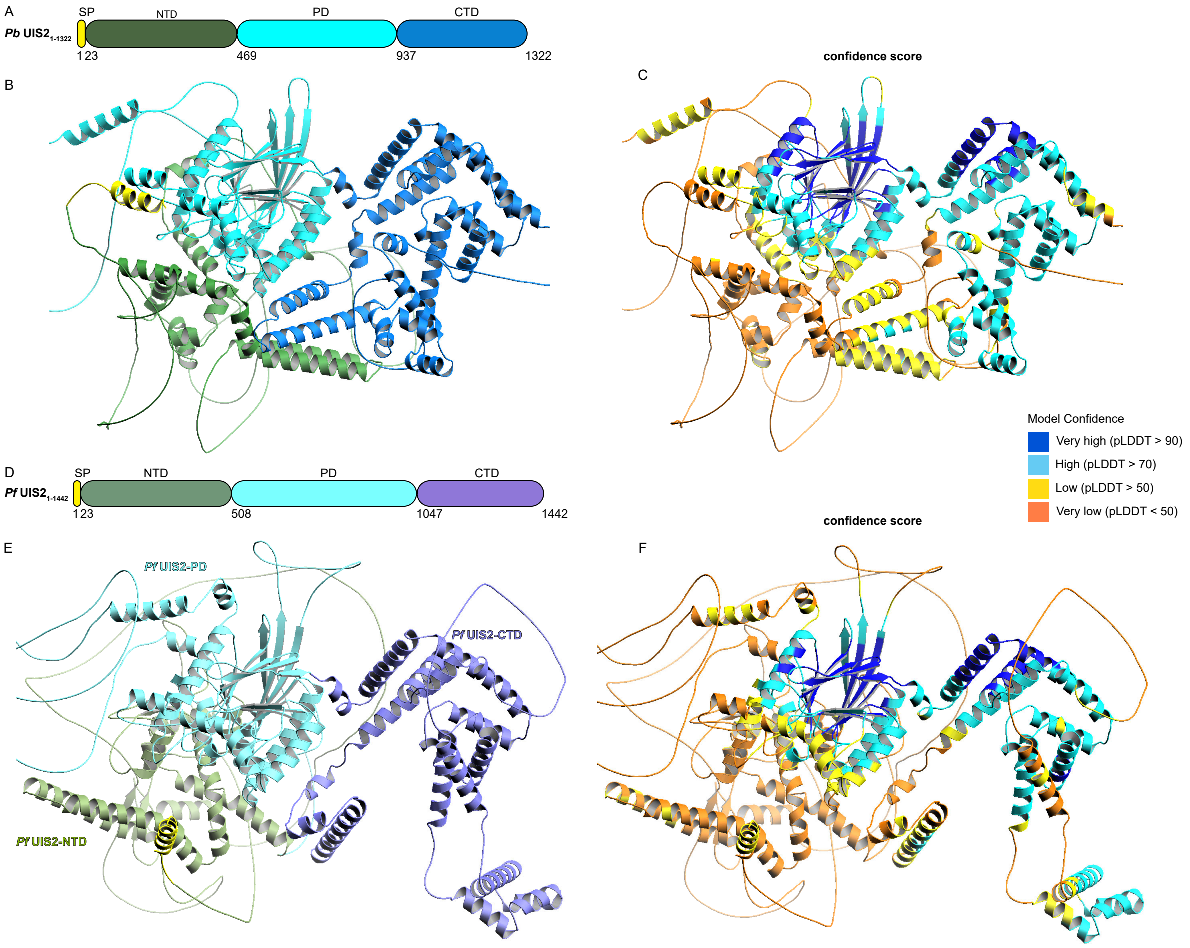


**Figure S3.** Alphafold predicts similar structures for UIS2 from *P. Berghei* and *P. Falciparum* with high confidence in the phosphatase domain. (*A*) Schematic diagram illustrating the domains within *Pb*UIS2: signal peptide (SP), N-terminal domain (NTD), phosphatase domain (PD), and C-terminal domain (CTD). (*B*) Predicted protein structure of *Pb*UIS2 (residues 1-1322) generated by AlphaFold2. (*C*) Predicted *Pb*UIS2 (residues 1-1322) structure color-coded based on the pLDDT confidence score. (*D*) Schematic diagram illustrating the domains within *Pf*UIS2 (residues 1-1442): signal peptide (SP), N-terminal domain (NTD), phosphatase domain (PD), and C-terminal domain (CTD). (*E*) Predicted protein structure of *Pf*UIS2 (residues 1-1442) generated by Alphafold2. (*F*) Predicted *Pf*UIS2 (residues 1-1442) structure color-coded based on the pLDDT confidence score.

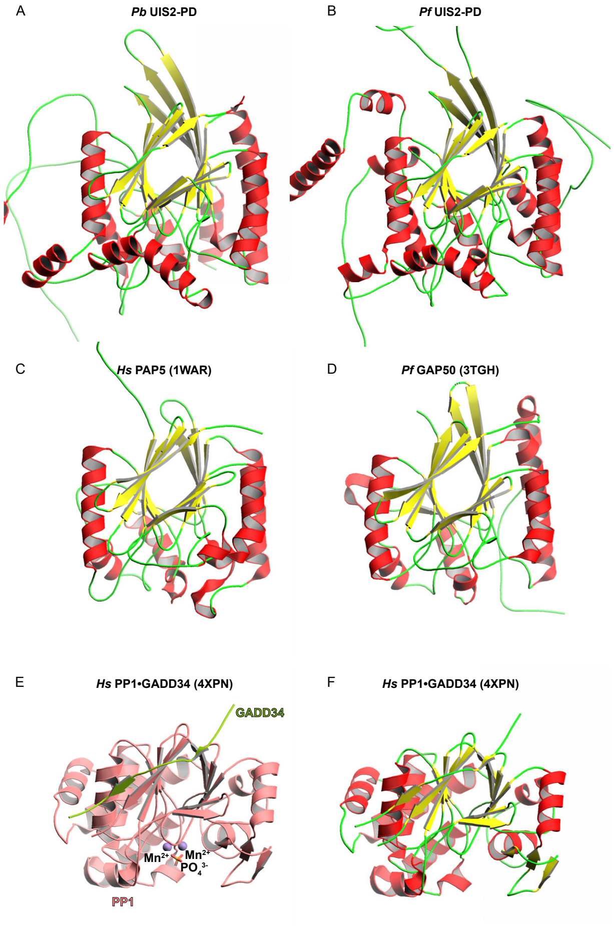


**Figure S4.** UIS2-PD exhibits structural similarities with PAP5 and GAP50 proteins rather than PP1. (*A-D*) Protein structures of the phosphatase domain in *Pb*UIS2 and *Pf*UIS2 predicted by Alphafold2, and human PAP5 and *P. falciparum* GAP50 obtained from the Protein Data Bank. Secondary structures are colored: $\alpha$-helices in red and $\beta$-strands in yellow. (*E-F*) Protein structure of GADD34:PP1 holoenzyme, obtained from the Protein Data Bank.

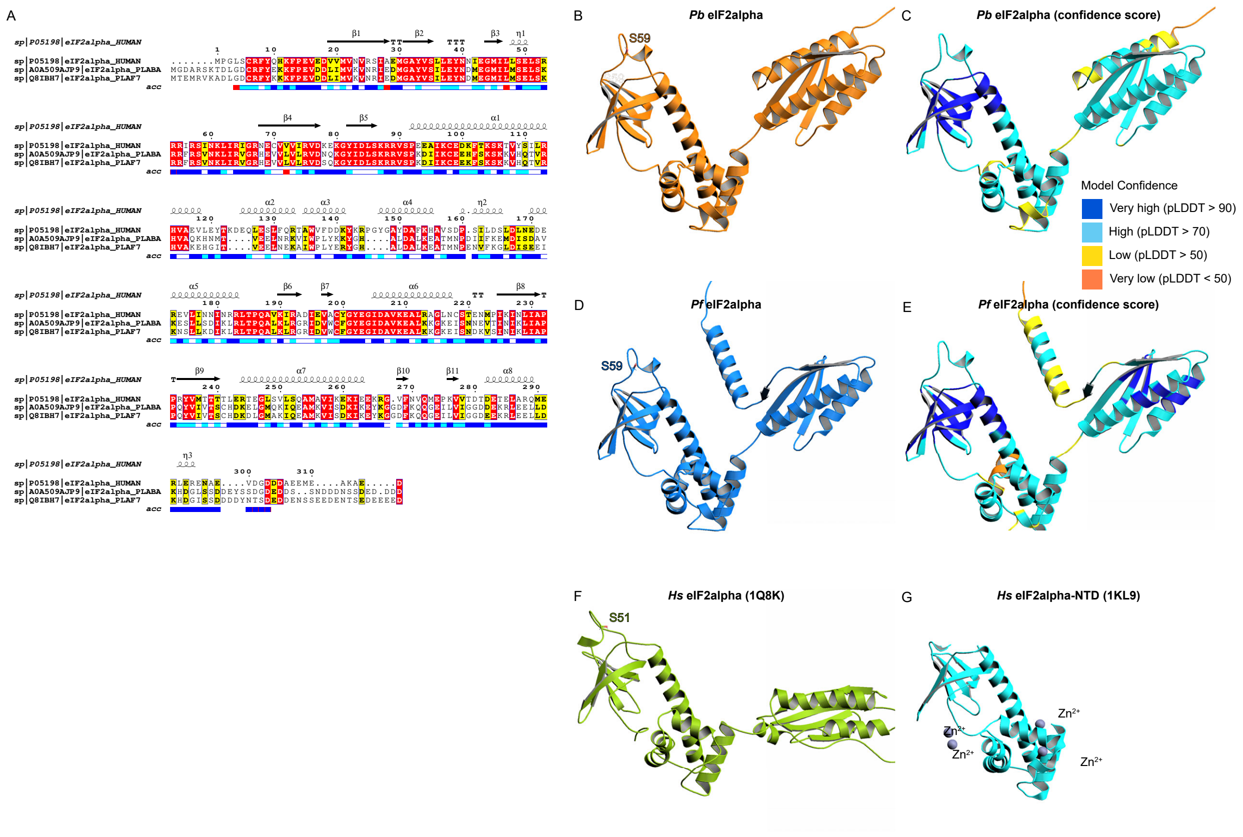


**Figure S5.** The eIF2alpha proteins from human, *P. berghei* and *P. falciparum* show sequence and structural homology. (*A*) Multiple sequence alignment of eIF2alpha protein sequences from human, *P. berghei*, and *P. falciparum*, performed using Clustal Omega and Escript 3.0. Secondary structure elements are indicated: $\alpha$-helices (squiggles), $3_{10}$-helices (small squiggles), $\beta$-strands (arrows), $\beta$-turns (TT letters). Conserved residues are highlighted in red within the sequence blocks. Accessibility is denoted by a bar below the sequences: blue for accessible, cyan for intermediate, and white for buried residues. (*B*) Predicted protein structure of *Pb*eIF2alpha generated by Alphafold2. (*C*) Predicted *Pb*UIS2 structure color-coded based on the pLDDT confidence score. (*D*) Predicted protein structure of *Pf*eIF2alpha generated by Alphafold2. (*E*) Predicted *Pf*eIF2alpha structure color-coded based on the pLDDT confidence score. (*F-G*) Protein structure of human eIF2alpha, obtained from the Protein Data Bank.

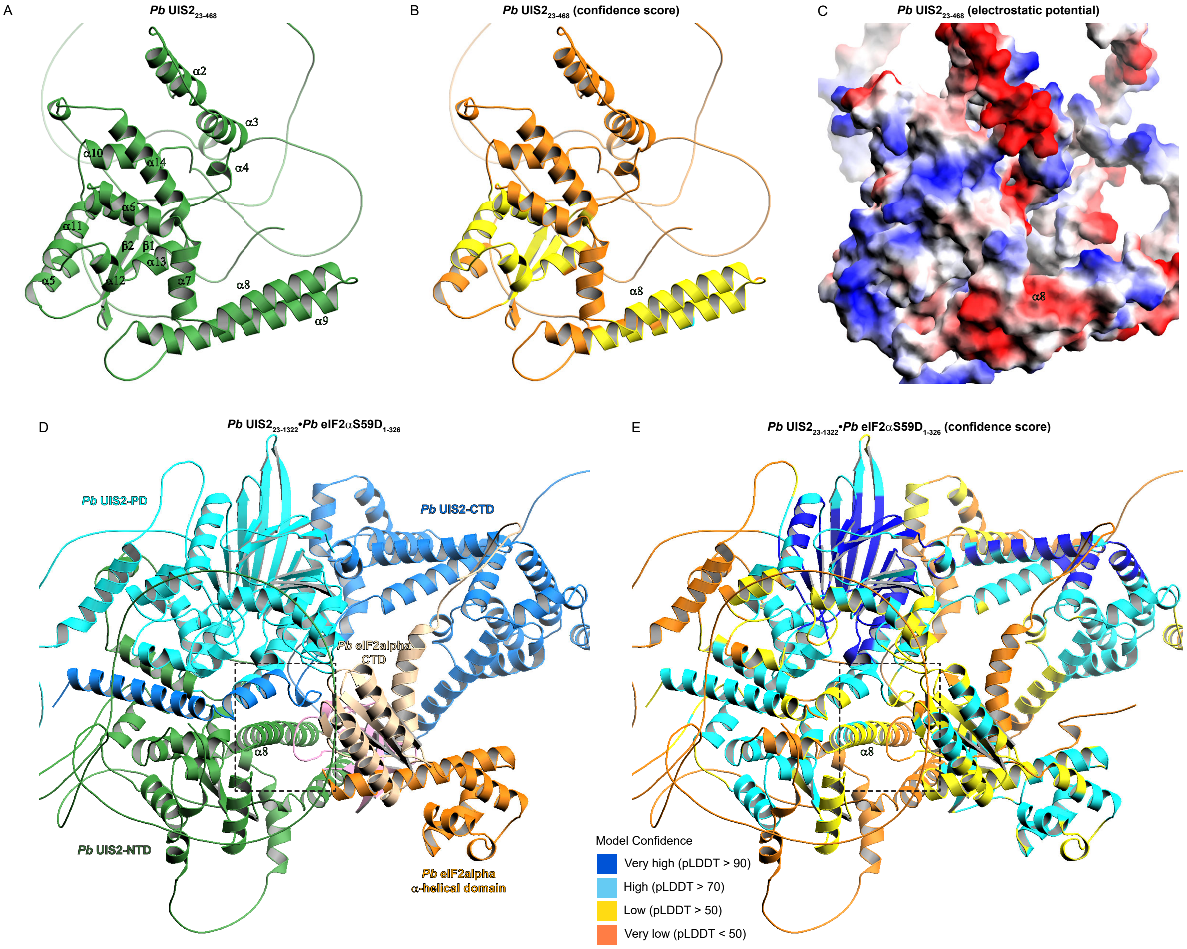


**Figure S6.** The loop region from the S1 domain in *Pb*eIF2alphaS59D interacts with α8 from *Pb*UIS2-NTD. (*A*) Predicted protein structure of *PbUIS2* (residues 23-468, NTD) generated by AlphaFold2. (*B*) Predicted *Pb*UIS2 (residues 23-468, NTD) structure color-coded based on the pLDDT confidence score. (*C*) *Pb*UIS2-NTD is displayed as an electrostatic surface representation, with red indicating negative charges and blue indicating positive charges. (*D*) Predicted protein complex structure formed by *Pb*UIS2 (residues 23-1322) and *Pb*eIF2alpha S59D, generated by the AlphaFold2 multimer approach. (*E*) Predicted *Pb*UIS2 (residues 23-1322) and *Pb*eIF2alpha S59D co-complex structure color-coded based on the pLDDT confidence score.

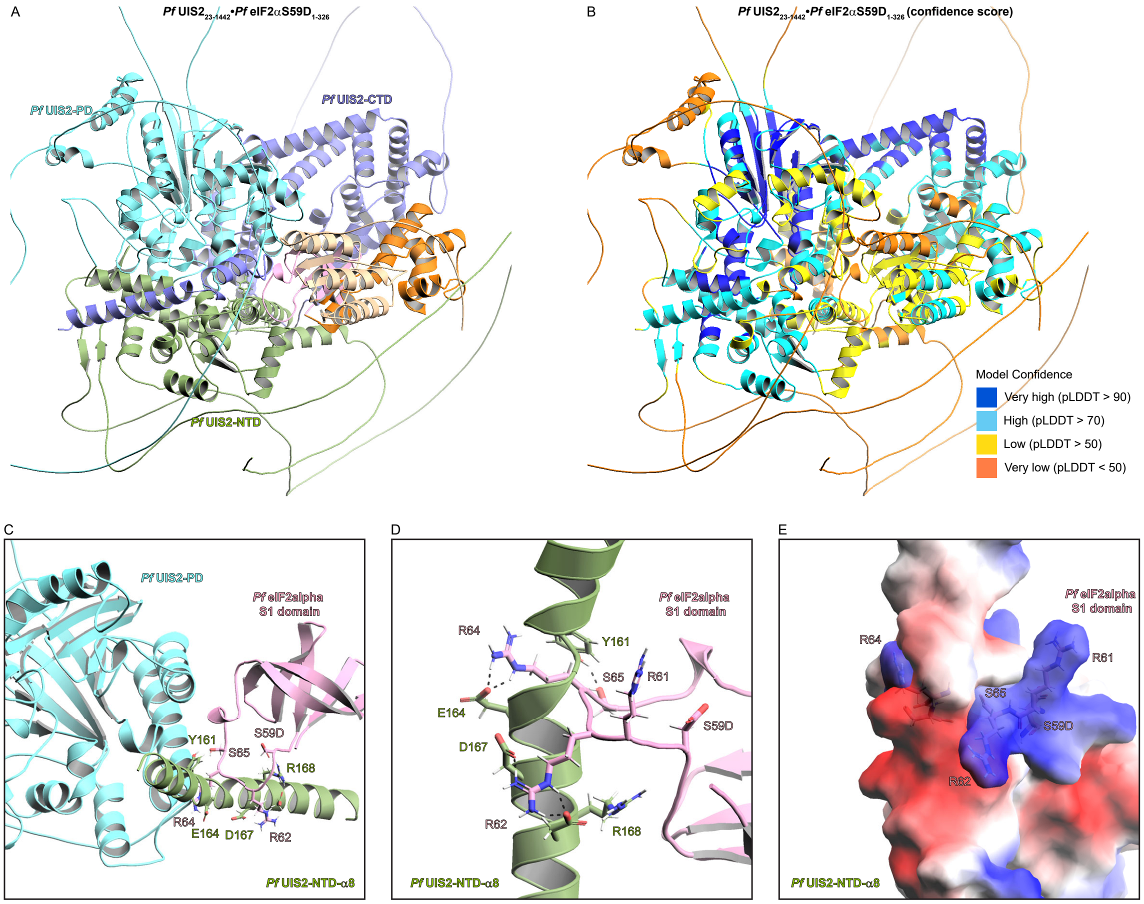


**Figure S7.** The loop region from the S1 domain in *Pf*eIF2alphaS59D interacts with α8 from *Pf*UIS2-NTD. (*A*) Predicted protein complex structure formed by *Pf*UIS2 (residues 23-1442) and *Pf*eIF2alpha S59D, generated by the AlphaFold2 multimer approach. (*B*) Predicted *Pf*UIS2 (residues 23-1442) and *Pf*eIF2alphaS59D co-complex structure color-coded based on the pLDDT confidence score. (*C*) Isolated view of the *Pf*UIS2 (residues 23-1442) and *Pf*eIF2alphaS59D complex from (*A*), showing only *Pf*UIS2's PD and helix 8 (α8) from the NTD, along with the S1 domain of *Pf*eIF2alphaS59D. Key residues involved in the protein interaction are depicted as sticks. (*D*) Close-up view of the interaction interface between *Pf*UIS2-NTD and the loop region from the S1 domain of *Pf*eIF2alphaS59D. Polar interactions are represented as black dashes. (*E*) The α8 from *Pf*UIS2-NTD and the loop region from the *Pb*eIF2alphaS59D S1 domain are displayed as an electrostatic surface representation, with red indicating negative charges and blue indicating positive charges.
